# Oscillating-flow Thermal Gradient PCR

**DOI:** 10.1101/544908

**Authors:** Varun L. Kopparthy, Niel D. Crews

## Abstract

We report the development of a versatile system based on oscillating-flow methodology in a thermal gradient system for nucleic acid analysis. Analysis of DNA and RNA samples were performed in the device, without additional temperature control and complexity. The technique reported in this study eliminates the need for predetermined fluidic channels for thermocycles, and complexity involved with additional incubation steps required for RNA amplification. A microfluidic device was fabricated using rapid prototyping by simply sandwiching dual side adhesive Kapton tape and a PDMS spacer between glass microscope slides. Amplification of the 181-bp segment of a viral phage DNA (ΦX174) and B2M gene in human RNA samples was demonstrated using the system. The developed system enables simultaneous acquisition of amplification and melt curves, eliminating the need for post-processing.

## INTRODUCTION

Worldwide, infectious diseases are the top ten causes of deaths^1^. Nucleic acid testing (NAT) enables the rapid and accurate detection of viruses and bacteria from a patient’s sample. Current NAT instruments are bulky, expensive, and slow, limiting the potential for point-of-care (POC) applications^2–4^. Lab-on-a-chip technology has led to the development of miniaturized NAT devices for POC applications. Polymerase chain reaction (PCR) and reverse transcription polymerase chain reaction (RT-PCR) are widely employed nucleic acid quantification techniques^5^. Amplification of the initial nucleic acid is achieved via PCR by thermocycling between temperature zones to perform denaturation (95°C), annealing (60°C), and extension (72°C). RT-PCR requires an additional incubation (45°C) step in which the RNA is first converted to complementary DNA (cDNA) by reverse transcriptase^6^. A multitude of PCR variants exist today that have advantageous and specific applications for nucleic acid analysis^5^. Several research groups have developed miniaturized PCR devices - based on sample thermocycling, they are broadly categorized into stationary chamber-based^7^, continuous-flow^8–10^, and oscillating-flow devices^11–14^. Existing miniaturized PCR devices require the consideration of a variety of parameters such as cycle number selection, number of thermal zones for various incubation steps, and multiple of single sample testing, before the device fabrication. A ‘one size fits all’ type is required to provide the user with the ability to determine the number of PCR cycles and add incubation steps required by the PCR type is required for widespread adaptation of the miniaturized PCR devices in POC applications.

Quantitative analysis of the nucleic acid in the sample is required for genetic testing. A combination of amplification and melt curve analysis is powerful for genotyping applications. Typically, an additional post processing step that takes about 30 minutes is required by existing NAT devices to obtain melt curves. Previously, we have reported the acquisition of amplification and melt curves using fluorescence images while thermocycling the sample^15–18^. This system was based on continuous flow in a thermal gradient, eliminating the need for stringent temperature control. Our earlier work was compatible for DNA analysis only, either an external incubation step or modification of the fluidic channel geometry is required for RNA amplification. Oscillating-flow method for nucleic acid amplification is gaining increasing attention due to simplicity in operation, multiplexing, and flexibility in implementing thermocycles. Existing oscillating-flow devices require offline detection of the amplified products such as gel electrophoresis and melt analysis. Thus, limiting the oscillating flow devices for quantitative use. We hypothesize that by combining an oscillating flow methodology with a thermal gradient system, a versatile nucleic acid analysis system can be realized for simultaneous acquisition of amplification and melt curves.

In this work, we have developed a microfluidic device with a simple channel geometry using rapid prototyping. Nucleic acid sample introduced into the fluidic channel via a syringe pump is oscillated in the device that is maintained under thermal gradient to enable temperatures required for PCR. A real-time acquisition of fluorescence images and analysis will simultaneously provide amplification and melt curves. By programming the syringe pump and heater control, the system can be expanded to perform a range of PCR procedures. To demonstrate the operation of the device, both DNA and RNA analysis were performed. Here, the design, development and characterization of proposed oscillatory-flow thermal gradient system is reported.

## Results and Discussion

The thermal gradient system (Fig. 1a) used in this study consists of a syringe pump that is programmed to oscillate the sample at a flow rate of 10 µl/min between denaturing and annealing temperatures in the microfluidic channel (Fig 2.) An in-house fabricated controller and heater system is used to generate thermal gradient across the microfluidic chip. The temperatures on the chip were maintained at 60°C and 90°C for denature and annealing. Using the optical system, fluorescence images were recorded during thermocycling to simultaneously obtain the amplification and melt curves (Fig. 1b).

**Figure 1:**
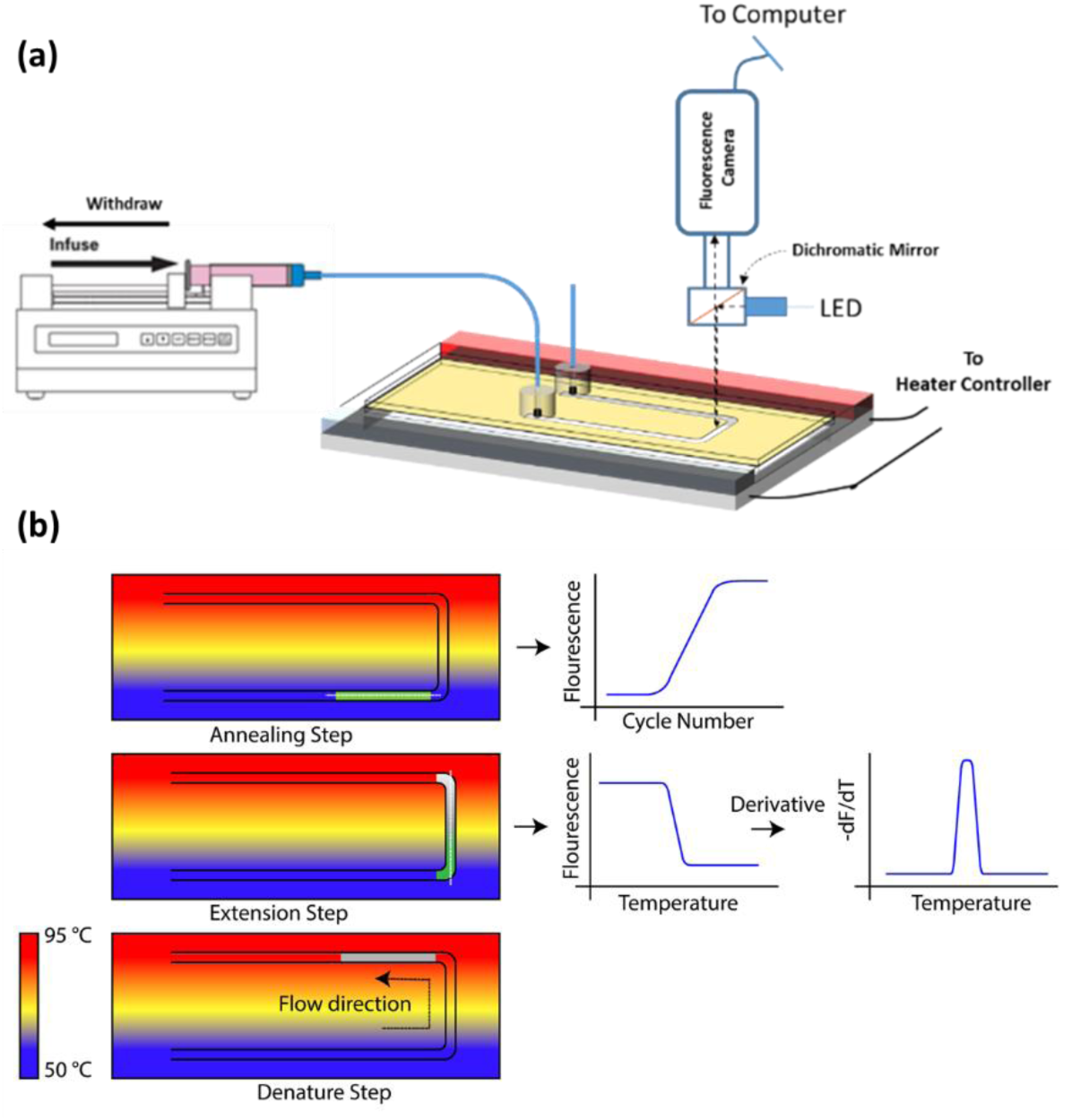
(a) Experimental setup of the thermal gradient oscillating-flow PCR system. (b) concept of thermal gradient system and image analysis for simultaneous amplification and melting curves.

**Figure 2:**
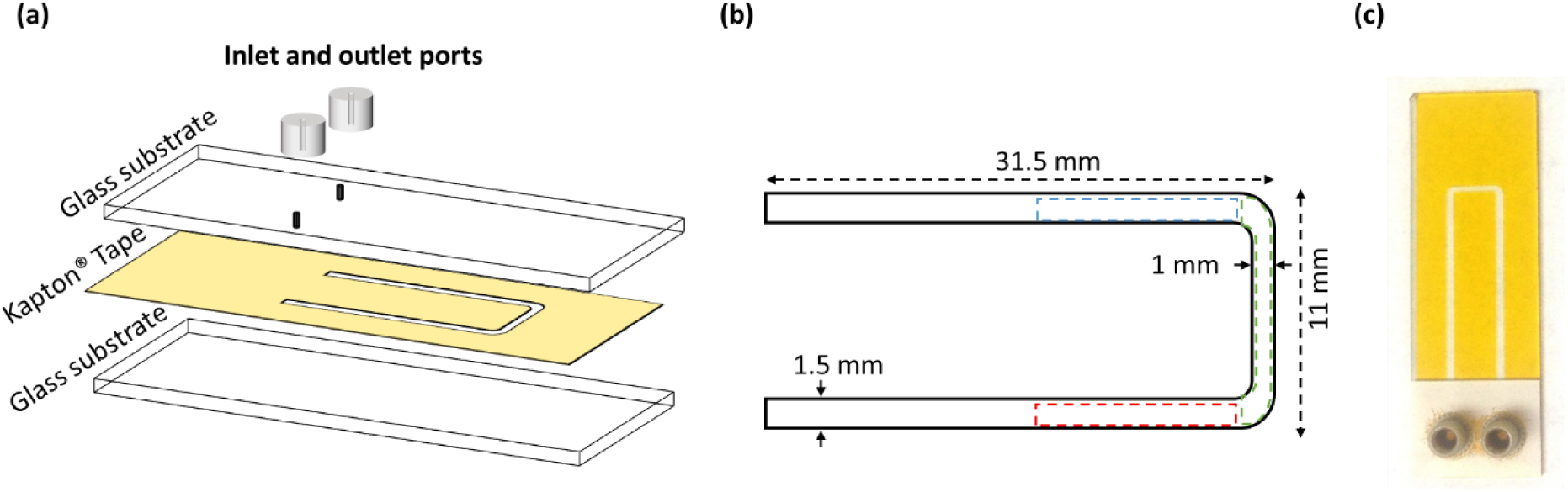
Schematic of the components of microfluidic device fabrication. (a) exploded view of microfluidic device showing glass slides, tape and inlet ports. (b) PCR channel dimensions showing annealing (blue dotted lines), extension (green dotted lines) and denaturation (red dotted lines) regions. (c) fabricated glass-tape composite microfluidic device.

### Thermal Calibration

The PCR is based on thermocycling of the sample; thermal performance of the device was characterized to determine the thermal variations. Infrared (IR) camera was used to analyze the thermal gradient on the chip. Isotherms on the device under no flow and flow conditions were evaluated. The device is loaded with a sample plug (water with 10% BSA) in Flourinert oil. The sample is oscillated between the cold and hot temperatures using the syringe pump. Infrared images of the temperature in the microfluidic chip under no flow, and flow conditions are shown in Figure 3(a, b, c). Temperatures can be extracted from the pixel values along the X-direction on the IR images. Three different lines in the Y direction are extracted and plotted to see temperature perturbations in all three conditions. Temperature values are plotted for comparison in Figure 3(d). The isotherms observed are straight when there is no flow in the microfluidic device. When the sample is oscillated between the annealing and denature temperatures, the flow causes a shift in the isotherms. Temperature perturbations of ≤ 1.5°C, and ≤ 3°C (Fig. 3e) are observed when the sample is moving from annealing to denaturing temperature, and from denaturing to annealing temperature, respectively (Fig. 3f).

**Figure 3:**
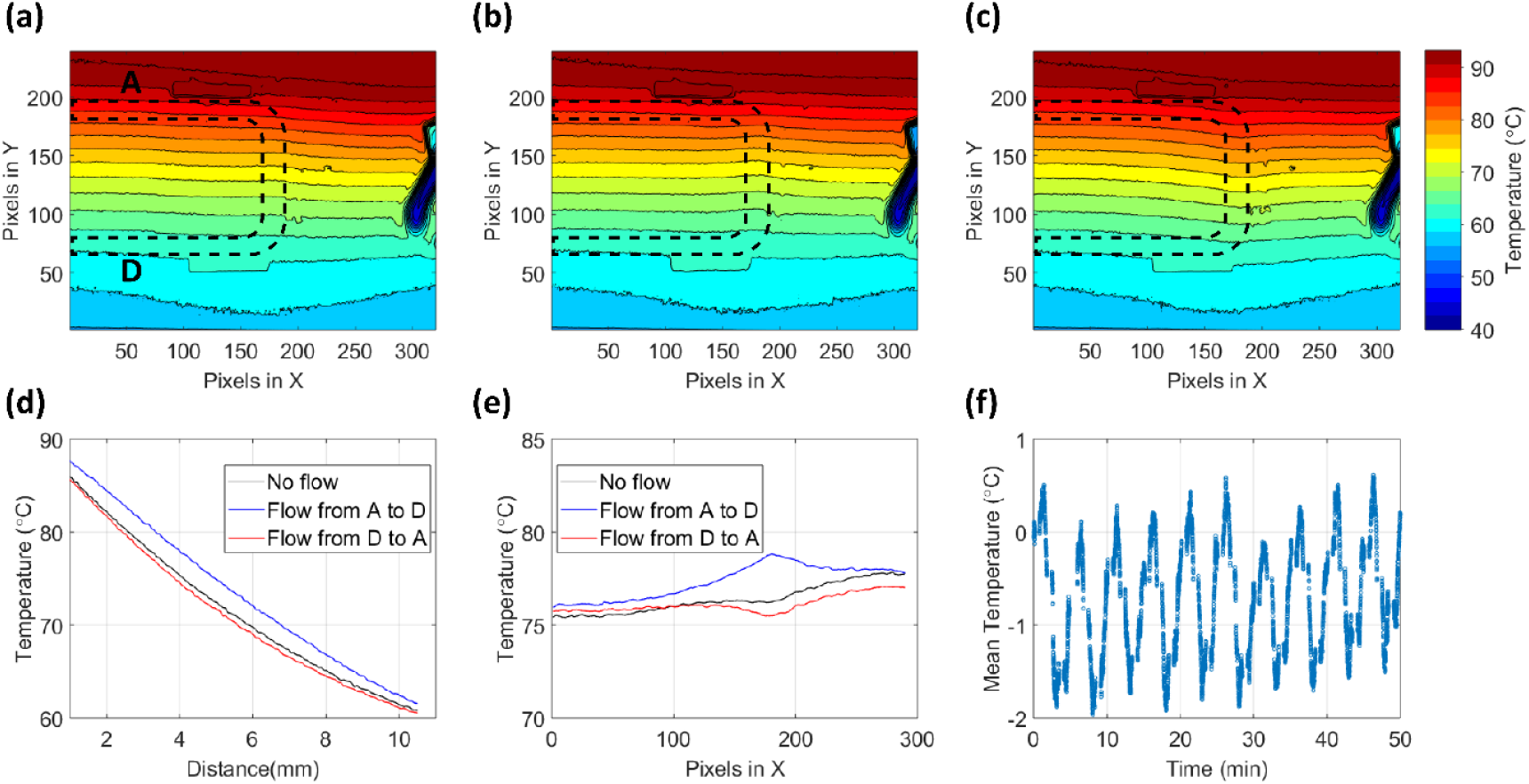
Infrared Images of the thermal gradient on the microfluidic device under (a) no-flow, (b) flow from annealing to denature temperatures, and (c) flow from denature to annealing temperatures. Channel geometry is shown in black dotted line. Arrow in white represents the direction of fluid flow. (d) Temperature perturbation plots extracted from IR images for no-flow, flow from annealing to denature temperatures, and flow from denature to annealing temperature conditions. (e) gradient variation under operating conditions. (f) temperature variation in the channel for a 40 cycle sample oscillation.

### Oscillating-flow PCR

While the PCR sample is oscillated in the microfluidic channel, fluorescence images (Fig. 4a) were recorded and analyzed to extract amplification and melt curves. Images recorded during the annealing stages provide the amplification curve, and images at the extension stage; when the sample is flown through the gradient, reveals the melting analysis. Increase in the fluorescence intensity is seen in the recorded images, indicating the amplification process. Extracting the fluorescence data and plotting against cycle number provides amplification curves. Figure 4b shows the standard melt curves extracted from the obtained fluorescence images. First order derivative of the standard melt produces melt peaks. Figure 5 compares data from the proposed microfluidic oscillating-flow PCR system and the commercially available Light Scanner (LS-32) system. The fluorescent image from the proposed system provides amplification and melting simultaneously. Unlike the LS-32 system, the oscillating-flow system provides the melting analysis at every stage, which is useful when multiple targets are amplified at the same time. Melting analysis provides the cycle at which the individual targets start to amplify.

**Figure 4:**
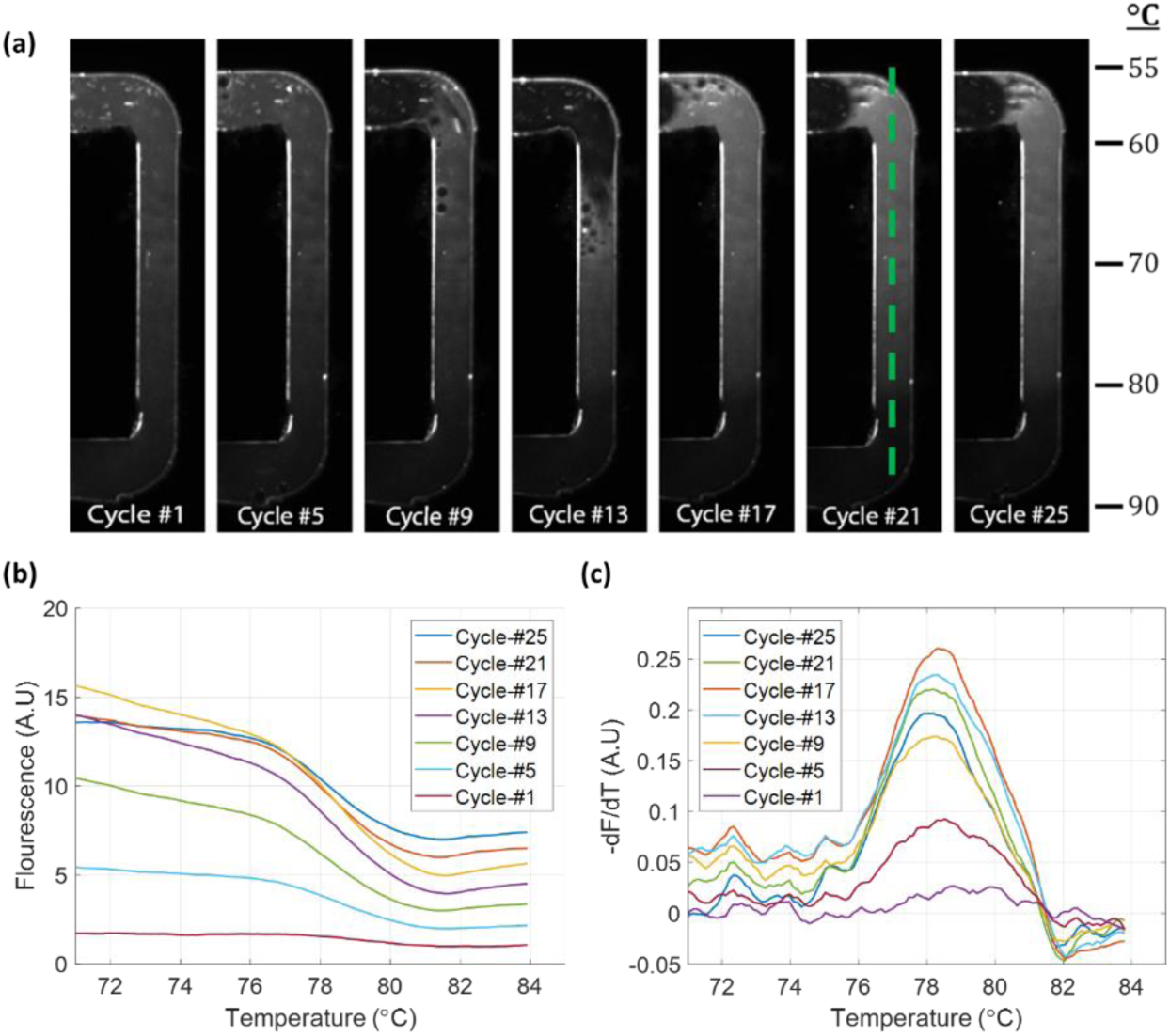
(a) Fluorescence images of melts at various cycles in the PCR. Analysis of the fluorescence in the channel along the dotted green line provides standard melt curves. (b) melt curves obtained from analyzing the fluorescence images. The melt curves are separated for better visualization. (c) melt peaks obtained by the first derivative of the standard melt curves.

**Figure 5:**
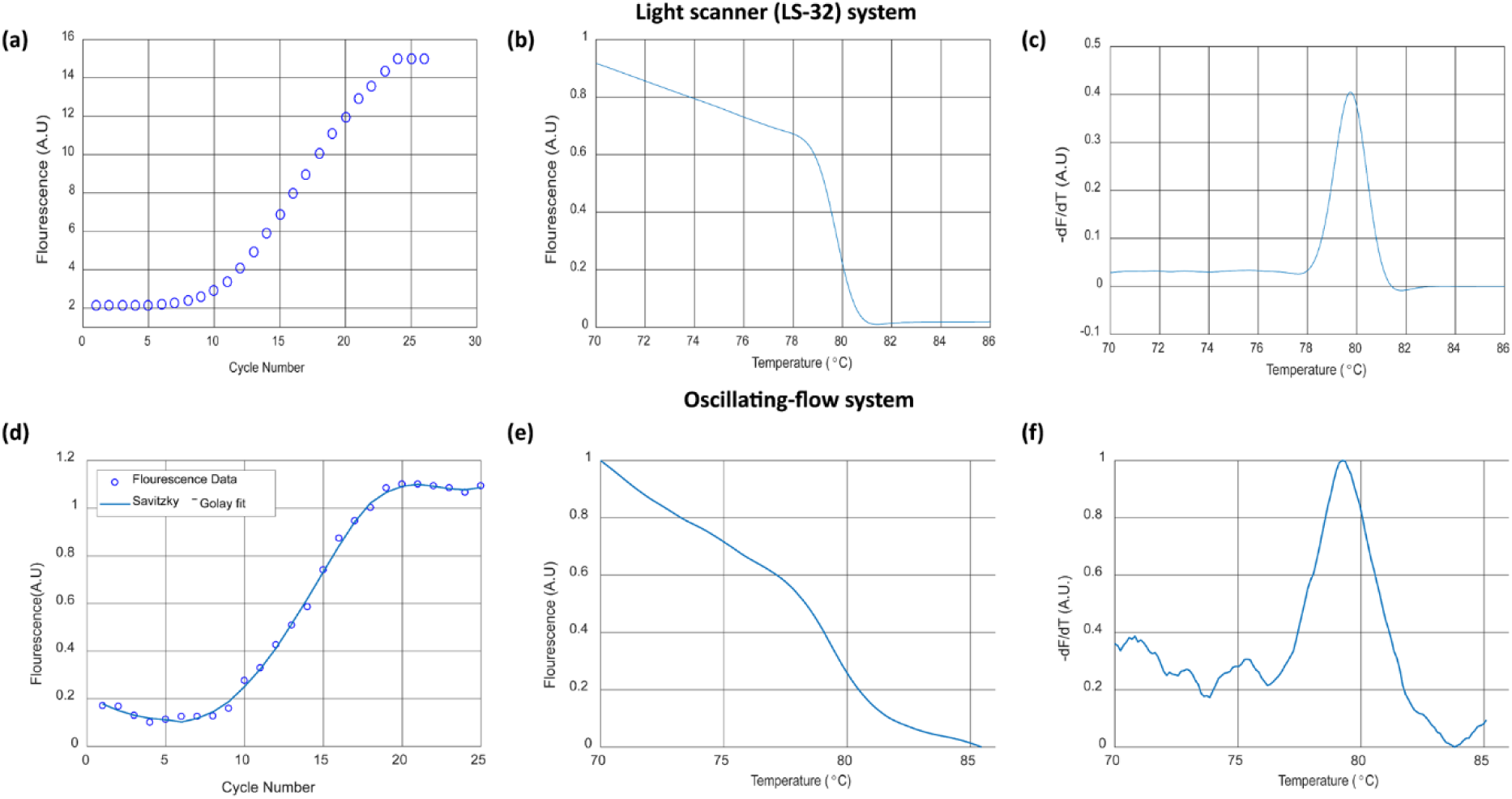
(a,b,c) are the amplification curve, standard melting curve, and melting peak of the phage DNA PCR using LS-32 system. (d,e,f) are the amplification curve, standard melt curve and the melting peak of the phage DNA PCR using the oscillating-flow system.

### On-chip RT-PCR

To perform RT-PCR, the PCR mix containing RNA sample is loaded into the microfluidic device and the temperature is raised isothermally to 50°C using the heater controller. After incubation for 5 minutes to reverse transcription of RNA to cDNA is achieved. The controller is then programmed with denature and annealing temperature set points to generate thermal gradient for performing traditional PCR. Up on thermocycling for 40 cycles, the sample is collected from the microfluidic device and a melt analysis (Fig. 6) is performed on the light scanner LS32 system. The melt data (Fig. 6b) shows the similar products are performed when amplified on-chip and in LS32 system. the melt peaks showed a slight variation for the product formed on the oscillating-flow device compared to LS32 system, possibly due to the variation in the dye and salt concentrations in the collected sample. The system was successfully utilized to perform RT-PCR of human RNA samples.

**Figure 6:**
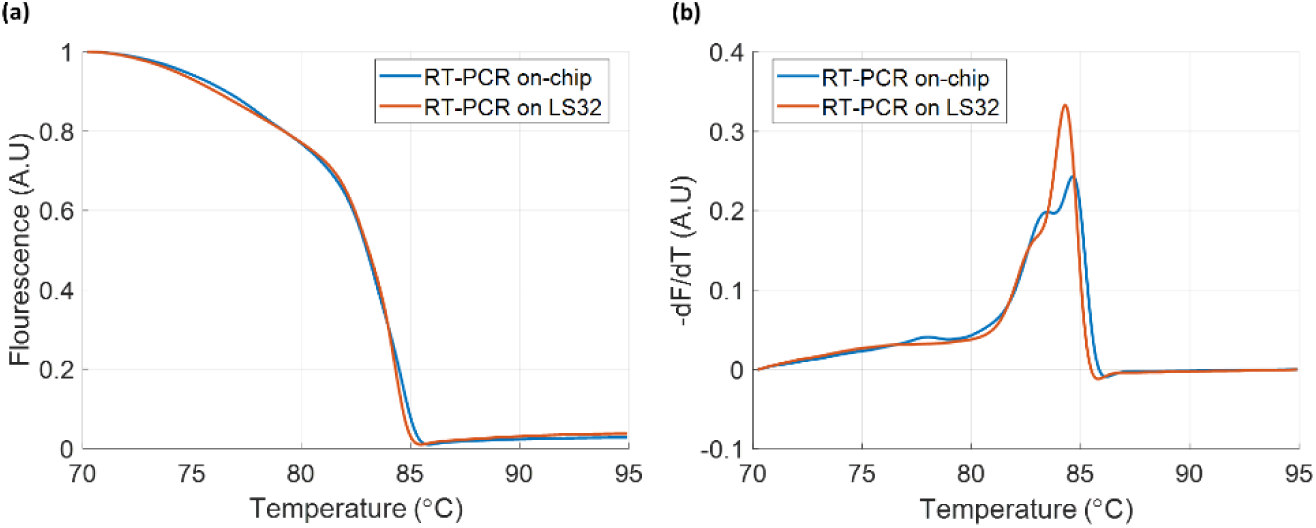
(a) High-resolution melt curves after RT-PCR in LS32 and in the developed oscillating-flow system. (b) First order derivatives of the high-resolution.

An oscillating-flow thermal gradient system for nucleic acid analysis is developed that can enable simultaneous amplification and melting analysis. The device operation was demonstrated by amplifying viral phage DNA and human RNA gene. Inexpensive and disposable microfluidic chips were fabricated using a rapid prototyping technique. The versatility of the system with the combination of a simple microfluidic device and programmable thermal gradient heater control and image acquisition - eliminates the need for predetermining cycle number selection, and post-PCR melt analysis. The oscillating-flow system demonstrated in this article showed excellent performance comparison with a standard benchtop real-time system. In the future, reducing the microfluidic channel dimensions will improve the PCR time and packaging the entire setup into a portable tool, would enable point-of-care nucleic acid analysis.

## MATERIALS AND METHODS

### Microfluidic device design and fabrication

To fabricate the microfluidic devices (figure 1), patterned double sided adhesive tape, a poly-dimethyl siloxane (PDMS) sheet, and a double sided adhesive polyimide tape were bonded between two aminosilane-coated glass slides (S4615, Sigma-Aldrich, MO, USA). A cutting plotter (Craft Robo Pro, Graphtech, USA) was used to pattern 100 µm double sided adhesive polyimide tape (PPTDE 1 112, Kaptontape.com, CA, USA) and a 250 µm thick PDMS sheet in the shape of the channel. A fine tip drill bit ((850-01OC, NTI, Kahla, Germany) was used to drill holes on a glass slide for fluid inlet and outlet in to the microchannel. The assembly of the microfluidic device with inlets is shown in Figure 1. Nanports (Upchurch scientific, WA, USA) were attached with superglue over the drilled holes. To clean the glass slide prior to the device fabrication, it was rinsed with 1% solution of detergent (Alconox, NY, USA), followed by distilled water, and dried with compressed air. PDMS sheet was rinsed with ethanol and dried with compressed air to improve bonding to the double-sided polyimide tape. Placing the device in a vice and tightening ensures strong bonding to the glass slides and reduces air bubbles when the device is subjected to heating during the experiment.

### PCR Reagents

For DNA amplification, the PCR mixture contained 108 copies/µl of a viral phage DNA template (ϕXI74, New England Biolabs, MA, USA), and 0.5 µM of each of the forward and reverse primers (Integrated DNA Technologies, IA, USA). Primers for 110 bp viral phage DNA target are F-GGTTCGTCAAGGACTGGTTT, R-TTGAACAGCATCGGACTCAG. For RNA amplification, human RNA (Integrated DNA Technologies, IA, USA), gene specific primer for B2M were used in the RT-PCR mixture. Reverse transcriptase enzyme (Takara Biosciences, USA) was used in one-step protocol for RT-PCR. Reverse transcription of RNA to cDNA requires an incubation at 42 °C for 5 min.

A PCR kit (Takara Biosciences, USA) was used to prepare the PCR mixture. A 2X one-step buffer containing deoxynucleotide triphosphate (dNTP), and hot start polymerase enzyme was used from the kit. 2.5 mg/ml bovine serum albumin (BSA) (Sigma-Aldrich MO, USA) was also used in the mixture. Intercalating dye, LC Green (LC Green Plus, Idaho Technology, UT, USA) was used to observe the fluorescence in PCR amplification in both the commercial system (Light scanner system LS-32, Idaho Technology, UT, USA) and the proposed oscillating-flow PCR system. The amplification protocol for the LS-32 consisted of a 1 min initial denaturation at 95 °C, followed by 30 cycles of 95 °C for 1 s, 60 °C for 1 s, and 75 °C for 3s. All temperature ramping on the LS-32 was at a rate of 5 °C/s. After the PCR, a high-resolution melting analysis of each amplified sample was performed serially by monitoring the fluorescence during a steady ramp of 0.3 °C/s from 60 °C to 90 °C.

### Microfluidic Chip Loading

The fabricated microfluidic chip is prepared for testing by flowing 100 µl of 1mg/mL BSA solution through the chip at a flow rate of 2 µl/min. BSA binds to the aminosilane channel walls and prevents the binding of the intercalating dye, LC Green. The chip was rinsed with DI water to remove excess BSA from the chip and is loaded with PCR mixture without any heating for passivating the channel surface. The chip is then emptied and loaded with the PCR sample with Fluorinert oil (FC40, Sigma Aldrich MO, USA) making the sample a droplet.

### Experimental

An infrared camera (A320, FLIR, OR, USA) with a 320×240-pixel array was used to calibrate and validate the temperatures and gradient on chip. A syringe pump (Pump Elite 11, Harvard Apparatus, USA) is programmed to introduce the PCR sample into the microfluidic device and oscillate between the annealing and denature regions. A monochrome 1392×1040 (1.4 MP) resolution camera (Pixel Link PL-B957U, ON, Canada) was used to capture images of the chip while the sample is oscillated between the denaturing and annealing temperatures. Blue LED lights, 470 nm (Luxeon Star LEDs, USA, MR-B0030-20T) were used as an excitation source for the fluorescence dye. A dichromatic mirror is used to allow excitation and emission in the same line of the camera. Filter sets were used to filter the LED wavelength and the emission wavelength to obtain a sharper quality image. The filter set and dichromatic mirror (Chroma Technology Corporation, USA) were selected based on the spectrum properties: LED filter 425-475 nm (HQ450/50x); camera filter 485-535 nm (HQ510/50nm). Adding a dichromatic mirror 380-750 nm (Q480LP) intensified the LED output on the microfluidic chip, improving the melting analysis image of the channels. Images were recorded for every 45 seconds with an exposure time of 1 second. Recorded images provide the PCR amplification curve and melting analysis simultaneously. Images were analyzed using MATLAB (The MathWorks, MA, USA) to obtain the amplification and melting curves.

### Data Availability

All data generated or analyzed during the current study are available from the corresponding author on reasonable request.

## Acknowledgements

N.C. and V.K. acknowledge primary funding support for this effort from the NASA under Grant LA 13-EPSCoR-0027 and in part by the NSF under Grant CBET 1151148.

## Author Contributions

N.C. and V.K conceived the experiments. V.K. conducted the experiments, performed data collection, and analyzed the results. N.C. and V.K. contributed to writing and editing the manuscript.

## Additional Information

Competing Interests: The authors declare no competing interests.

